# Heterologous expression of carbonic anhydrase in *Acinetobacter* sp. Tol 5 for whole-cell biocatalysis

**DOI:** 10.1101/2025.06.09.658560

**Authors:** Shogo Yoshimoto, Hiroya Oka, Yuki Ohara, Yan-Yu Chen, Masahito Ishikawa, Katsutoshi Hori

**Author notes:** Corresponding author: Katsutoshi Hori. US Department of Energy Joint Genome Institute, Lawrence Berkeley National Laboratory, Berkeley, CA 94720, USA.

## Abstract

Carbonic anhydrase accelerates the hydration of carbon dioxide (CO_2_) and is an attractive biocatalyst for carbon capture and utilization. *Acinetobacter* sp. Tol 5 shows high adhesiveness via its cell-surface protein AtaA. We previously demonstrated its application to bacterial immobilization and gas-phase bioproduction. Here, we developed Tol 5 cells expressing carbonic anhydrase and evaluated CO_2_ conversion ability as whole-cell biocatalysts. A codon-optimized carbonic anhydrase from *Sulfurihydrogenibium yellowstonense* (SyCA) was produced in the cytoplasm, but the cells showed little activity as a whole-cell biocatalyst. To enhance activity, we fused six signal peptides (SPs) to SyCA for periplasmic expression. The Omp38-SP fusion of SyCA was properly processed to the mature size, yielding higher whole-cell activity. By contrast, the other constructs were either undetectable or remained unprocessed, resulting in lower activities. These results show that periplasmic expression of SyCA is important for efficient CO_2_ hydration in Tol 5 cells as whole-cell biocatalysts.

## Introduction

The amount of carbon dioxide (CO_2_) emissions as a result of human activity increased from 24.9 to 41 billion tons per year in the period from 1990 to 2022 emphasizing the urgent need for efficient, scalable carbon capture and utilization technologies (Yurak and Fedorov, 2025). A variety of strategies, including physical absorption, chemical solvent scrubbing, membrane separation, and physical and chemical adsorption, have been explored to mitigate CO_2_ emissions (Hanifa et al., 2023; Kammerer et al., 2023; Nagireddi et al., 2023). Among them, biocatalytic and bio-integrated approaches that operate under ambient temperature and pressure without organic solvents are methods currently attracting significant attention because of their sustainability (Carceller et al., 2023; Marra et al., 2024; Sieborg et al., 2024).

Carbonic anhydrase (EC 4.2.1.1) is a zinc-dependent metalloenzyme that accelerates the reversible hydration of CO_2_ to bicarbonate (HCO_3_^-^). Solubilized bicarbonate can be captured as calcium carbonate or utilized by microorganisms as a metabolic substrate (Liu et al., 2020; Talekar et al., 2022). The catalytic efficiency of carbonic anhydrase is very high, with a *k*_*cat*_ value of 10^6^ s^-1^, thereby enhancing CO_2_ solubility by several orders of magnitude (Khalifah, 1971; Talekar et al., 2022; Shao et al., 2024). Many studies have shown that purified carbonic anhydrase markedly accelerates CO_2_ capture (Talekar et al., 2022); however, the scale required for industrial carbon capture makes the use of purified enzymes economically impractical.

Consequently, whole-cell catalysts that do not require a costly purification process are gaining attention as a cost-effective alternative (Moon et al., 2020; Ting et al., 2022; Zhu et al., 2022; Baidya et al., 2024).

The gram-negative bacterium *Acinetobacter* sp. Tol 5 exhibits high adhesiveness to various material surfaces, including hydrophobic plastics, hydrophilic glass, and metals, through its fibrous cell surface protein AtaA (Ishikawa et al., 2012). We previously developed a method for immobilizing bacterial cells using AtaA; bacterial cells transformed with the *ataA* gene exhibit nonspecific high adhesiveness, and large amounts of growing and resting cells can be quickly and firmly immobilized onto various material surfaces (Ishikawa et al., 2014; Hori et al., 2015). The immobilized cells directly on surfaces through AtaA show better tolerance and increased chemical reaction rates and can be repeatedly used in reactions without inactivation (Ishikawa et al., 2014). Furthermore, cells immobilized via AtaA can be placed directly in the gas phase, enabling efficient reactions with gaseous substrates (Yoshimoto et al., 2017; Usami et al., 2020). Therefore, Tol 5 is expected to be useful for CO_2_ utilization. However, to the best of our knowledge, there have been no reports of expressing a heterologous carbonic anhydrase in *Acinetobacter*.

In this study, we aimed to develop Tol 5 expressing carbonic anhydrase and to evaluate CO_2_ conversion ability as a whole-cell biocatalyst.

## Materials and methods Bacterial strains

The restriction-modification system and *ataA*-deficient strain of *Acinetobacter* sp. Tol 5 REK123Δ*ataA* (Tol 5_REK_) was used as the host strain (Ishikawa and Hori, 2024; Inoue et al., 2025). *Escherichia coli* DH5α was used for plasmid construction. Tol 5_REK_ was cultured in LB medium at 28°C, and DH5α was cultured in LB medium at 37°C.

### Plasmid construction

The oligonucleotides and synthetic DNA used in this study are listed in Table S1. An *Acinetobacter* codon-optimized gene encoding SyCA (carbonic anhydrase from *S. yellowstonense*) amino acid residues 21–246 was synthesized with a ribosome-binding site immediately upstream and a C-terminal hexahistidine tag followed by a stop codon and the T7 terminator. The synthesized gene was inserted into the EcoRI and XbaI sites of pARP3, generating pARP3-SyCA. To construct a plasmid expressing SyCA fused to a signal peptide (SP), the SyCA coding region was amplified by PCR from pARP3-SyCA using primers SyCA-F and SyCA-R, producing Fragment-1. For PelB, TorA, and Omp38, complementary oligonucleotides (SP-F and SP-R) were annealed to yield the double-stranded fragment (Fragment-2). For Tat-1, Tat-2, or Tat-3, complementary oligonucleotides (SP-F1 and SP-R1) were annealed, and a DNA fragment was amplified by PCR using primers SP-F2 and SP-R2 with the double-stranded oligonucleotides as a template, yielding Fragment-2′. The backbone of pARP3-SyCA was linearized with EcoRI and XbaI to obtain Fragment-3. Fragments 1, 2 (or 2′), and 3 were assembled using HiFi DNA Assembly Master Mix (New England Biolabs, Ipswich, United Kingdom), generating the expression plasmid for SP-fused SyCA. The constructed plasmid was introduced into Tol 5_REK_ cells by electroporation as previously described(Ishikawa and Hori, 2024).

### Protein expression

One milliliter of the overnight culture of Tol 5_REK_ harboring the expression plasmid was inoculated into 100 ml of LB medium supplemented with gentamicin (10 µg/mL), 0.5% arabinose, and 0.5 mM zinc sulfate and cultured at 28°C for 18 h at 115 rpm. Cells were harvested by centrifugation at 5,000 × g for 5 min at 4°C. The cell pellet was resuspended in SDS-sample buffer (5% (v/v) 2-mercaptoethanol, 2% (w/v) SDS, 0.02% (w/v) bromophenol blue, 62.5 mM Tris-HCl, pH 6.8) and subjected to SDS-PAGE followed by western blotting using an anti-His-tag antibody to verify expression.

### Wilbur-Anderson unit (WAU) assay

CO_2_ hydration activity was evaluated using the Wilbur and Anderson unit (WAU) assay (Wilbur and Anderson, 1948) with slight modifications. Cultured cells were harvested by centrifugation at 5,000 × g for 5 min at 4°C, and the cell pellet was resuspended in pure water at an optical density of 1.0 at 660 nm. Eight milliliters of 20 mM Tris-HCl (pH 8.3) and 1 ml of the cell suspension were added to a 30-ml beaker on ice.

Immediately after 6 ml of chilled CO_2_-saturated water was added to the beaker, the pH drop was recorded from 8.3 to 6.3 with stirring on ice using a pH sensor (PH-208; SATOTECH, Kanagawa, Japan) equipped with a logger (MJ-LOG2; SATOTEC). The obtained times for the cell suspension sample and blank (water) were designated T and T_0_, respectively. The WAU was defined as (T_0_ - T)/T. Differences among the samples were evaluated using the Brunner–Munzel test with Holm’s step-down adjustment.

## Results

### Cytoplasmic expression of SyCA

The α-type carbonic anhydrase from *Sulfurihydrogenibium yellowstonense* (SyCA) (Capasso et al., 2012) was chosen as the model enzyme because of its high catalytic efficiency and remarkable thermostability. A synthetic SyCA gene, codon-optimized for *Acinetobacter*, was cloned into the *Acinetobacter* expression plasmid pARP3 under the control of the arabinose-inducible promoter, with a C-terminal His-tag for detection.

The constructed plasmid was introduced into Tol 5_REK_, a restriction-modification system and *ataA*-deficient strain of Tol 5, for accurate enzymatic activity measurement due to its non-agglutination nature. SDS-PAGE of whole cell samples followed by western blotting with an anti-His antibody revealed a clear band corresponding to SyCA, confirming successful cytoplasmic expression (Fig. 1A).

**Figure 1.**
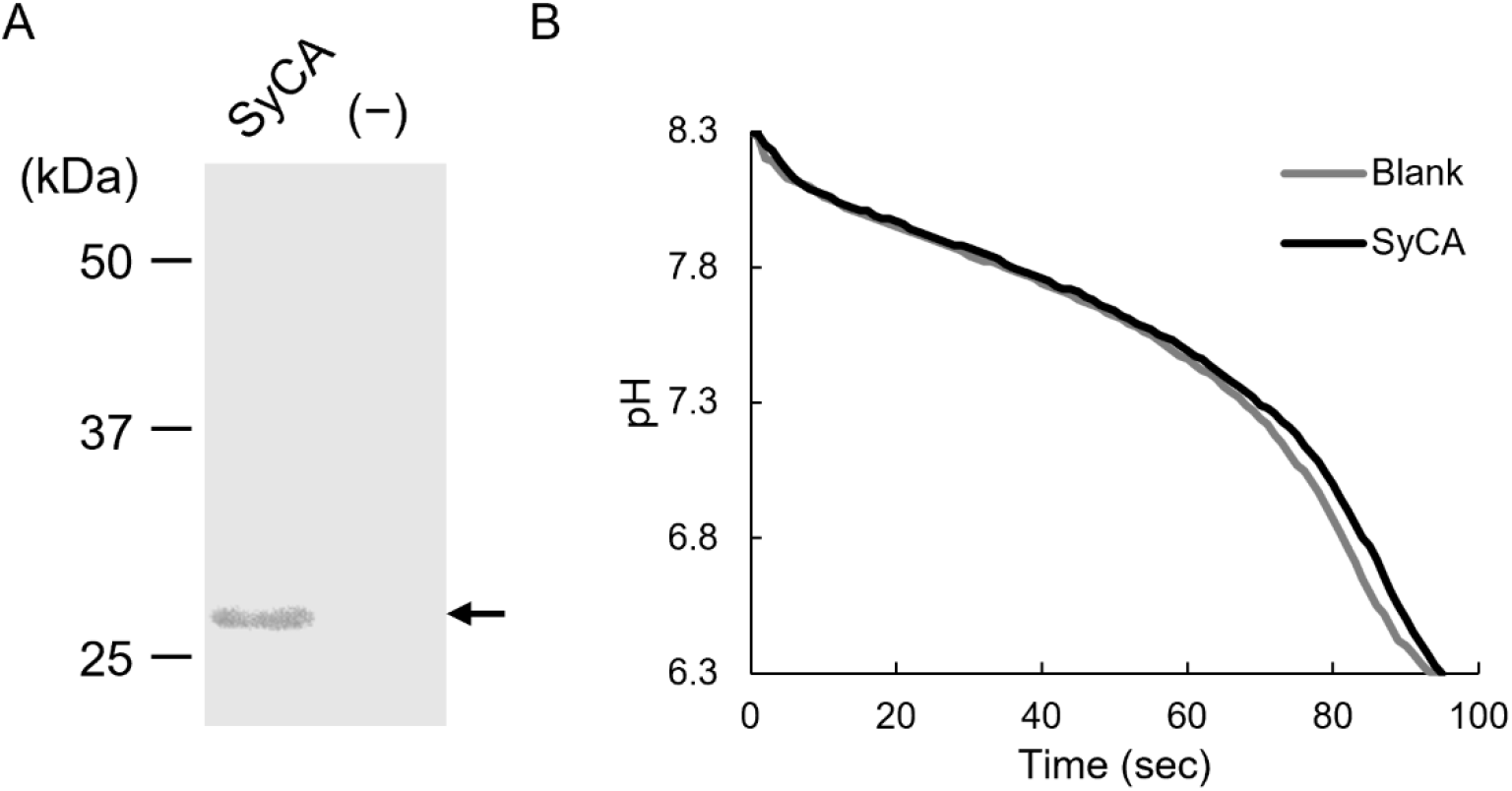
Cytoplasmic expression of SyCA. (A) Western blotting of whole-cell lysates from Tol 5_REK_ cells carrying the SyCA expression plasmid (+) or carrying no plasmid (–). Proteins were separated by SDS-PAGE, transferred to a PVDF membrane, and probed with an anti-His antibody; the arrowhead marks the band corresponding to SyCA. (B) Time course of the pH changes during the WAU assay. Carbonic anhydrase activity of cells expressing cytoplasmic SyCA or blank (water only, no cells) was measured.

Whole-cell carbonic anhydrase activity was assessed with the Wilbur– Anderson unit (WAU) assay (Wilbur and Anderson, 1948), which measures the rate of CO_2_ hydration to bicarbonate via the associated pH drop in CO_2_-saturated water. A representative pH–time profile is shown in Fig. 1B. Cells expressing cytoplasmic SyCA produced a similar pH–time curve as the blank, which was water instead of cell suspension, and the time required to reach pH 6.3 was almost the same. These results indicate that the cytoplasmic SyCA exhibited little activity as a whole-cell biocatalyst under the conditions.

### Selection of signal peptides for the periplasmic expression of SyCA

A previous study has shown that cytoplasmic expression of carbonic anhydrase leads to poor enzymatic activity because the cell membrane limits CO_2_ access and can even cause intracellular toxicity, presumably via pH imbalance (Jo et al., 2013). Therefore, periplasmic expression is regarded as an alternative strategy for efficient heterologous expression of carbonic anhydrase. We thus attempted to express SyCA in the periplasm of Tol 5_REK_.

Proteins that need to be secreted to the periplasm are mediated by either the general secretion (Sec) pathway or the twin-arginine translocation (Tat) pathway (Mergulhao et al., 2005; Natale et al., 2008). In the Sec pathway, the unfolded polypeptide is secreted from the cytoplasm to the periplasm through the SecYEG channel. A signal peptidase then cuts off its N-terminal signal peptide (SP), and the protein folds in the periplasm. In the Tat system, polypeptides that have already folded are secreted from the cytoplasm to the periplasm through the TatABC complex, and then their SPs are cleaved by a signal peptidase. To identify suitable signal peptides (SPs) for exporting SyCA, we first selected two well-established *E. coli* SPs(Matos et al., 2012; Jo et al., 2013): PelB for the Sec pathway and TorA for the twin-arginine translocation (Tat) pathway, as candidates. The Sec-type SP of Omp38 (Ishikawa et al., 2016), which is an abundant OM protein known as OmpA (Confer and Ayalew, 2013), was also selected as a native Sec-type SP from Tol 5. Because no Tat-type SPs have been characterized in Tol 5, open reading frames of the Tol 5 genome were screened with SignalP (Teufel et al., 2022), yielding three non-lipoproteins (BCX74450, BCX75992, and BCX75163), two lipoproteins (BCX75681 and BCX73201), and one protein of uncertain classification (BCX72454). We selected one representative from each of the three groups possessing putative Tat-type SPs, resulting in three candidates: BCX72454, BCX75163, and BCX75681, which we named Tat-1, Tat-2, and Tat-3, respectively. In total, six SPs were selected (Table 1). Each SP sequence, spanning from the start methionine to its predicted cleavage site, was fused to the N-terminus of SyCA through a short triglycine (GGG) linker (Fig. 2A). The six fusion genes were cloned into the pARP3 vector, with a C-terminal His-tag, and were introduced into Tol 5_REK_.

**Table 1.**
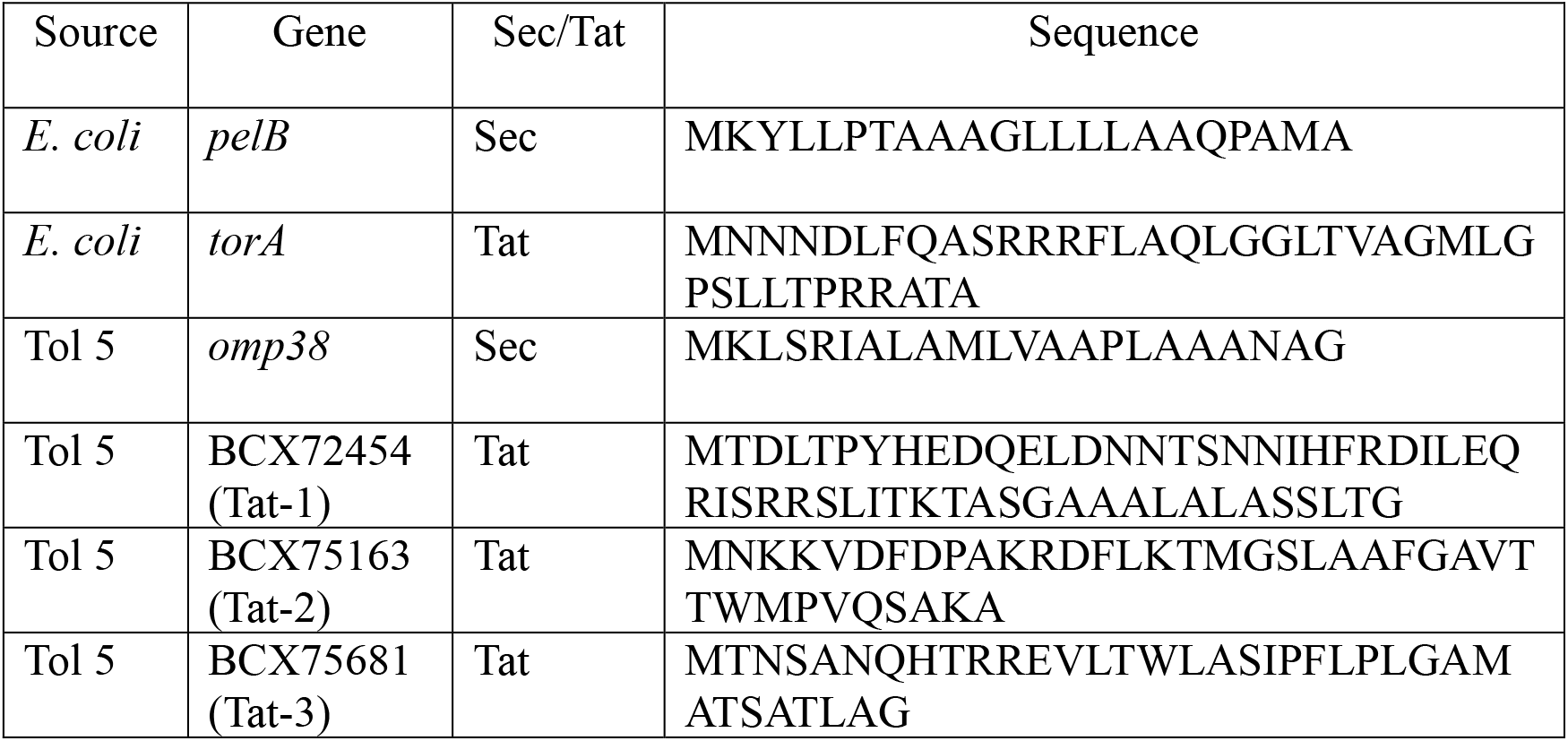
Signal peptides used in this study.

**Figure 2.**
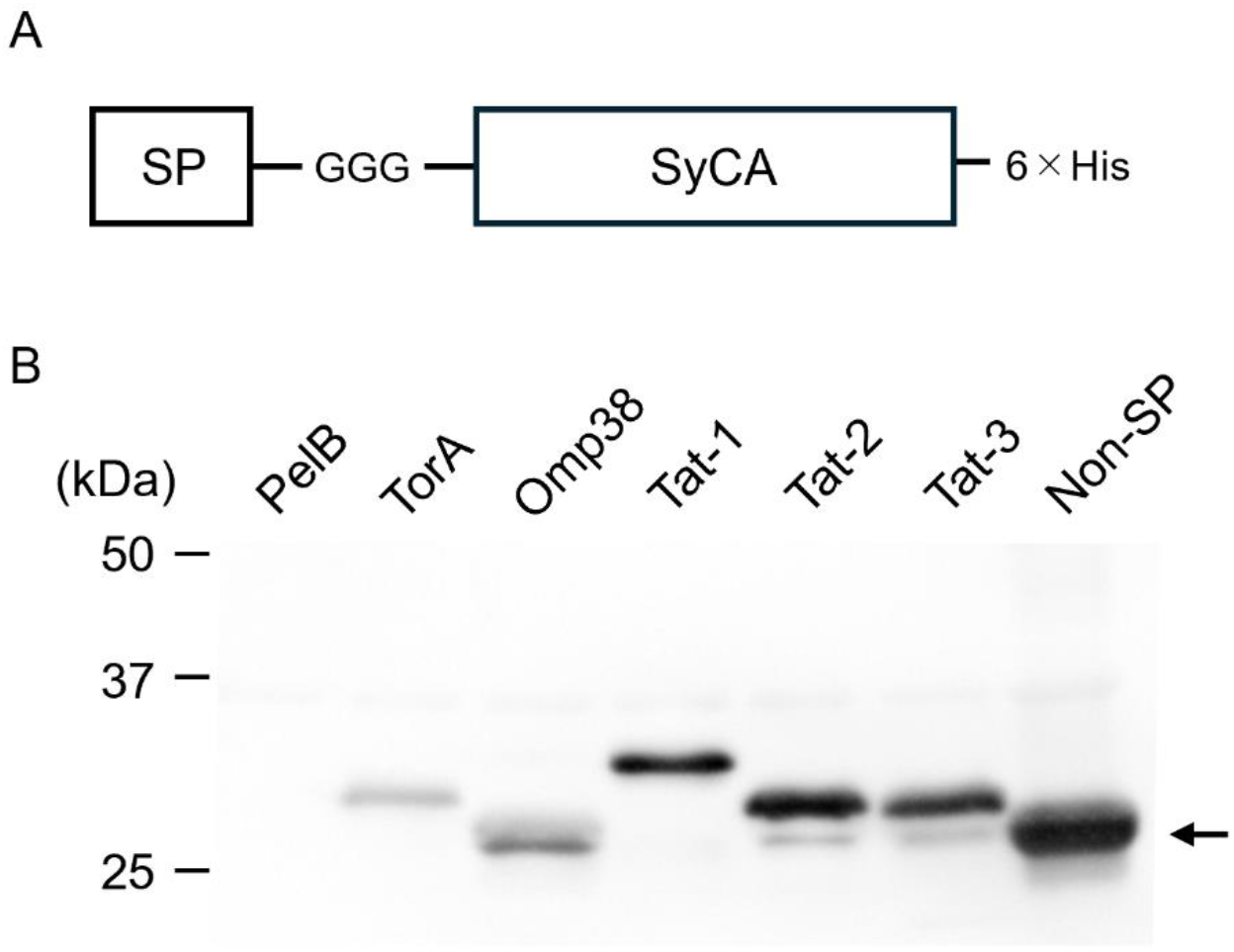
Expression of SyCA variants fused to each signal sequence. (A) Schematic representation of SyCA fused to a signal peptide (SP). (B) Protein expression was detected by western blotting using an anti-His antibody. The arrowhead indicates the band corresponding to the cytoplasmically expressed SyCA (Non-SP).

### Expression of SyCA fused to each signal peptide

Tol 5_REK_ cells harboring the expression plasmid were cultured at 28 °C for 18 h with 0.5 % arabinose to induce the expression of SyCA. After cell lysis, proteins were analyzed by western blotting with an anti-His antibody. SyCA was detected in every construct except when fused to the SP of PelB (Fig. 2B). The constructs fused to SPs from TorA, Tat-1, Tat-2, and Tat-3 exhibited slightly higher molecular weights than the cytoplasmically expressed SyCA, suggesting that their SPs were not removed. In contrast, the SyCA fused to the SP of Omp38 showed a band at the same size as the cytoplasmic SyCA, indicating proper cleavage of its SP and secretion into the periplasm. These results suggest that the Omp38-SP is the most suitable for periplasmic expression of SyCA in Tol 5_REK_.

### Catalytic activity of cells expressing SyCA fused to each signal peptide

We evaluated whether fusion with the SPs improved whole-cell carbonic anhydrase activity. All SyCA-producing constructs, except the one fused to PelB-SP, were tested using the WAU assay. Figure 3A shows the representative pH-time profile: cells expressing SyCA fused to each SP lowered the pH faster than the blank, which was water instead of a cell suspension. This suggests that these cells exhibit carbonic anhydrase activity as a whole-cell biocatalyst. The WAU values calculated from three independent measurements were summarized in Figure 3B. The Omp38-SP, which follows the Sec pathway, yielded the highest activity. Tat-type SPs from either *E. coli* (TorA) or Tol 5 gave lower activity. The Brunner–Munzel test was applied with Omp38 as the reference. After Holm adjustment for four comparisons, Omp38 showed significantly higher WAU than TorA (adjusted *p* < 0.01), Tat-2 (adjusted *p* < 0.01), Tat-3 (adjusted *p* < 0.01), and Tat-1 (adjusted *p* = 0.013). These results indicate that periplasmic expression with a suitable SP is important for achieving effective whole-cell biocatalysis of carbonic anhydrase in Tol 5.

**Figure 3.**
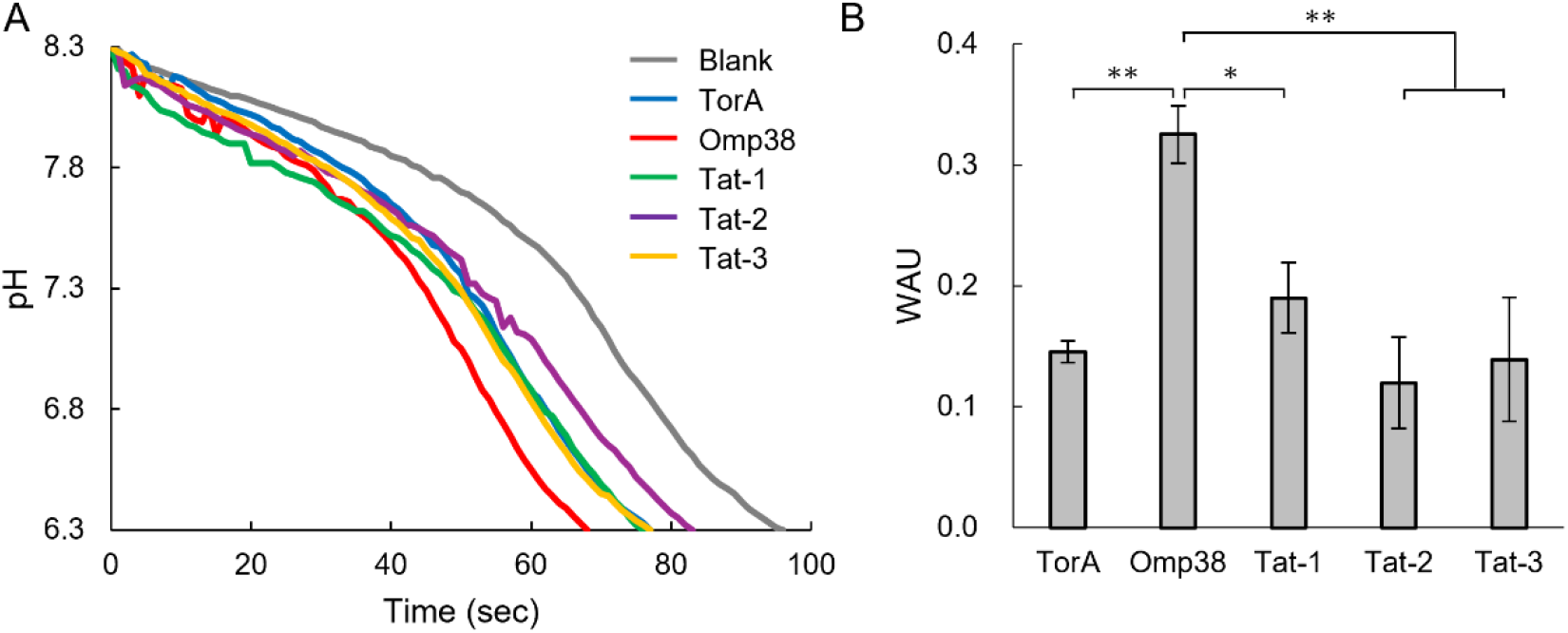
Carbonic anhydrase activity of cells expressing SyCA variants fused to each signal sequence. (A) Time course of the pH changes during the WAU assay of cells expressing SyCA variants fused to each signal sequence. (B) The WAU values calculated from three independent measurements. Data are expressed as mean ± SEM (n = 3). Statistical significance, *adjusted *p* = 0.013, **adjusted *p* < 0.01.

## Discussion

In this study, we investigated the expression of the carbonic anhydrase SyCA in Tol 5_REK_ and its CO_2_ conversion activity as a whole-cell biocatalyst. Fusing SyCA with the SP of Tol 5 Omp38 led to efficient processing of its SP, resulting in the highest WAU value (Figs. 2, 3). These results highlight that periplasmic expression and the selection of an appropriate SP are important for the catalytic performance of carbonic anhydrase as a whole-cell biocatalyst. To the best of our knowledge, this is the first report on the heterologous expression of carbonic anhydrase in *Acinetobacter* for carbon utilization.

Several studies have explored the periplasmic expression of carbonic anhydrase in *E. coli*. Jo *et al*. expressed NgCA using the PelB (Sec) and TorA (Tat) signal peptides and found that the TorA construct yielded higher catalytic activity (Jo et al., 2013). Patel *et al*. expressed the Cab and Cam with PelB (Sec) and gIII (Sec) signals, reporting that PelB conferred slightly higher activity for both enzymes (Patel et al., 2013). More recently, Jo *et al*. showed that the periplasmic expression of TdCA with the PelB (Sec) signal peptide enhanced enzyme activity (Jo and Hwang, 2020). Collectively, these findings indicate that the optimal signal peptide for periplasmic expression appears to be specific to each carbonic anhydrase even in *E. coli*. In Tol 5, the *E. coli* PelB-SP did not yield detectable SyCA production (Fig. 2), and the *E. coli* TorA-SP produced only weak activity (Fig. 3). These results suggest that SPs from *E. coli* do not match the secretion machinery of Tol 5 and are consistent with previous reports that variations in signal peptide sequences influence secretion efficiency and that some signal peptides exhibit species specificity (Blaudeck et al., 2001; Freudl, 2018). The three Tat-type SPs from Tol 5 were not fully cleaved and also yielded lower activity (Figs. 2, 3). In contrast, the native Sec-type Omp38-SP from Tol 5 was cleaved efficiently and gave the best activity, demonstrating its suitability for expressing SyCA in this host.

Tol 5 cells can be immobilized on many materials through AtaA and can be applied to gas-phase reactions (Yoshimoto et al., 2017; Usami et al., 2020). Combining the bacterial cell immobilization with the periplasmic expression of SyCA may lead to stable, reusable catalysts for gas-phase CO_2_ hydration or for supplying bicarbonate to downstream biosynthetic steps. Such systems could contribute to efficient biological carbon capture and conversion technologies.

## Conclusion

In this study, we demonstrated that periplasmic expression and selection of an appropriate SP are important for the catalytic performance of carbonic anhydrase in Tol 5 as a whole-cell biocatalyst. Integrating the adhesive properties of Tol 5 with engineered periplasmic expression of SyCA could lead to efficient CO_2_ capture and conversion processes.

## Author contribution

**Shogo Yoshimoto**: Conceptualization, Validation, Writing - Original Draft. **Hiroya Oka**: Investigation. **Yuki Ohara**: Investigation. **Yan-Yu Chen**: Investigation. **Masahito Ishikawa**: Investigation. **Katsutoshi Hori**: Conceptualization, Writing - Review & Editing, Funding Acquisition

## Acknowledgements and Funding information

The authors thank Kensho Abe, Chiharu Kitano, and Eriko Kawamoto for their technical assistance. An expression plasmid for SyCA used in preliminary experiments was kindly provided by I-Son Ng (National Cheng Kung University). This research was supported by the Japan Society for the Promotion of Science (JSPS) KAKENHI (Grant Number JP24H00043) to KH and SY.

## Data availability

All data generated or analyzed during this study are included in this published article and its supplementary materials.

