## Supplementary material for "Heterologous expression of carbonic anhydrase in *Acinetobacter* sp. Tol 5 for whole-cell biocatalysis": Table S1

**Table S1. Oligo nucleotides and synthetic DNA used in this study.**

| Name | Sequence |
| --- | --- |
| SyCA-F | GGAGGTGGAGAACATGAATGGTCGTATGAAGG |
| SyCA-R | GCCGCTCTAGAATCCGGATATAG |
| SP-PelB-F | GGGCTAGCGAATTCAAAGAGGAGAAAGGATCTATGAAATACCTGCTGCCGACCGCTGCTGCTGGTCTGCTGCTCCTC |
| SP-PelB-R | CATACGACCATTCATGTTCTCCACCTCCGGCCATCGCCGGCTGGGCAGCGAGGAGCAGCAGACCAGCAGCAGCGGTCGGCAGCAGGTATTTCATAGATCCTTTCTCCTCTTTGAATTCGCTAGCCCAAA |
| SP-TorA-F | GGGCTAGCGAATTCAAAGAGGAGAAAGGATCTATGAACAATAACGATCTCTTTCAGGCATCACGTCGGCGTTTTCTGGCACAACTCGGCGGCTTAACCGTCGCCGGGATGC |
| SP-TorA-R | CCTTCATACGACCATTCATGTTCTCCACCTCCCGCAGTCGCACGTCGCGGCGTTAACAATGACGGCCCCAGCATCCCGGCGACGGTTAAG |
| SP-Omp38-F | GGGCTAGCGAATTCAAAGAGGAGAAAGGATCTATGAAATTGAGTCGTATCGCACTTGCTATGCTTGTTGCTGCTCCAC |
| SP-Omp38-R | CCTTCATACGACCATTCATGTTCTCCACCTCCACCAGCATTAGCAGCAGCGAGTGGAGCAGCAACAAGCATAG |
| SP-Tat1-F1 | GATCAAGAGTTAGATAACAATACTTCGAATAACATTCATTTCCGTGATATTTTAGAACAACGGATTTCTCGTCGTAGTTT |
| SP-Tat1-F2 | GGGCTAGCGAATTCAAAGAGGAGAAAGGATCTATGACAGACCTCACGCCATATCATGAAGATCAAGAGTTAGATAACAAT |
| SP-Tat1-R1 | AAACTACGACGAGAAATCCGTTGTTCTAAAATATCACGGAAATGAATGTTATTCGAAGTATTGTTATCTAACTCTTGATC |
| SP-Tat1-R2 | CCTTCATACGACCATTCATGTTCTCCACCTCCACCTGTCAAGCTAGATGCTAAAGCTAATGCTGCTGCACCACTGGCTGTTTTGGTAATTAAACTACGACGAGAAATCCGTGCAGCGGTCTCCTCATCTTGTTTTCTACGCCTTTGATTTTGC |
| SP-Tat2-F1 | GATTTCGATCCAGCAAAACGTGATTTTTTAAAAACAATGGGAAGCCTTGCTGCATTTGGTGCCGTGACTACATGGATGCC |
| SP-Tat2-F2 | GGGCTAGCGAATTCAAAGAGGAGAAAGGATCTATGAATAAGAAAGTTGATTTCGATCCAGCAAAACGTG |
| SP-Tat2-R1 | GGCATCCATGTAGTCACGGCACCAAATGCAGCAAGGCTTCCCATTGTTTTTAAAAAATCACGTTTTGCTGGATCGAAATC |
| SP-Tat2-R2 | CCTTCATACGACCATTCATGTTCTCCACCTCCAGCCTTGGCACTTTGAACAGGCATCCATGTAGTCACGGC |
| SP-Tat3-F1 | GTCGTGAAGTGCTTACATGGTTAGCCAGCATTCCGTTTCTTCCACTTGGGGC |
| SP-Tat3-F2 | GGGCTAGCGAATTCAAAGAGGAGAAAGGATCTATGACAAATTCAGCCAACCAACACACACGTCGTGAAGTGCTTACATG |
| SP-Tat3-R1 | GCCCCAAGTGGAAGAAACGGAATGCTGGCTAACCATGTAAGCACTTCACGAC |
| SP-Tat3-R2 | CCTTCATACGACCATTCATGTTCTCCACCTCCACCTGCAAGAGTAGCAGAAGTTGCCATTGCCCCAAGTGGAAGAAACGG |
| Synthetic DNA | GAATTCAAAGAGGAGAAAGGATCTATGAAATTGAGTCGTATCGCACTTGCTATGCTTGTTGCTGCTCCACTCGCTGCTGCTAATGCTGGTGGAGGTGGAGAACATGAATGGTCGTATGAAGGTGAAAAAGGACCTGAACATTGGGCTCAATTAAAACCGGAATTTTTCTGGTGCAAACTCAAAAACCAATCTCCGATCAATATAGACAAGAAGTACAAAGTGAAAGCCAATTTACCTAAGCTTAACTTGTATTACAAAACAGCAAAAGAATCCGAAGTGGTGAATAATGGACACACTATTCAGATCAACATCAAAGAGGATAATACCCTAAATTATCTTGGAGAGAAATACCAGTTGAAGCAATTCCACTTTCATACTCCTTCTGAACATACGATAGAAAAGAAAAGCTATCCATTGGAAATTCATTTTGTCCACAAAACCGAAGATGGCAAAATCCTTGTTGTTGGTGTTATGGCGAAATTAGGGAAGACGAATAAGGAACTTGACAAAATACTCAACGTTGCTCCTGCAGAAGAAGGCGAAAAAATTCTGGATAAAAACTTGAACTTAAACAATCTGATTCCAAAAGATAAACGCTATATGACCTATAGTGGCTCATTGACAACTCCACCATGTACAGAGGGTGTTAGGTGGATTGTACTGAAAAAGCCCATAAGCATATCCAAACAACAACTTGAGAAGTTAAAGTCGGTCATGGTCAATCCCAATAATCGTCCAGTACAGGAGATTAATTCACGATGGATCATTGAGGGTTTTTTGGAGCATCATCATCACCATCACTAAGATCCGGCTGCTAACAAAGCCCGAAAGGAAGCTGAGTTGGCTGCTGCACCGCTGACAATAACTAGCATAACCCCTTGGGGCCTCTAAACGGGTCTTGAGGGGTTTTTTGCTGAAAGGAGGAACTATATCCGGATTCTAGA |
